# Screening, Characterization and Optimization of antibacterial peptides, produced by *Bacillus safensis* strain MK-12 isolated from waste dump soil KP, Pakistan

**DOI:** 10.1101/308205

**Authors:** Sajid Iqbal, Muhammad Qasim, Farida Begum, Hazir Rahman, Imran Sajid

**Affiliations:** Department of Microbiology, Kohat University of Science and Technology, Kohat KP Pakistan/ National University of sciences and Technology, Islamabad Pakistan; Department of Microbiology, Kohat University of Science and Technology, Kohat KP Pakistan; Department of Biochemistry, Abdul Wali Khan University Mardan, KP Pakistan; Department of Microbiology, Abdul Wali Khan University Mardan, KP Pakistan; Department of Microbiology and Molecular Genetics (MMG), University of the Punjab, Lahore Pakistan

**Keywords:** Soil bacteria, Peptides, Antibacterial activity, Optimization, Multidrug resistant

## Abstract

**Aims:** The current study was designed to isolate, screen and identify the indigenous soil antibacterial exhibiting bacteria (AEB) and effect of various parameters on growth of AEB and antibacterial peptides production.

**Methods and results:** The soil isolates were screened for antagonistic activity against a set of ATCC and local MDR human pathogenic bacterial strains. The antibacterial compound was protein in nature, exhibited no haemolysis and molecular weight was less than 20 KDa. The potential AEB isolate was identified by morphology, biochemical testing and by 16S rRNA gene sequencing as *B. safensis* MK-12. Growth and antibacterial activity was optimized for *B. safensis* strain MK-12, exhibited maximum growth as well as antibacterial activity after 48 hours of incubation at pH 8, 30 °C in shaking incubator when fermented in optimized medium.

**Conclusion:** The current study results indicate that indigenous soil is rich source of AEB and could be a promising source of antimicrobial compounds to fight against MDR bacteria in future.

**Significance and impact:** This is the first scientific report on soil bacteria from northern region of Pakistan as per our knowledge. Therefore, screening of soil bacteria for antibacterial activity from unexplored area may contribute towards new antibiotic. Selected soil strain in the current study exhibited promising antibacterial activity against human pathogenic MDR bacterial strains.

## Introduction

In the mid of 20th century, when antibiotics were yet not available to the patients. The individuals with bacteremia or septicemia had low chances of survival. The availability of antibacterial drugs and synthesis of new chemotherapy’s agents reversed this scenario. Recently, the emergence of multi drug resistant (MDR) in pathogenic bacterial strains such as *S.aureus* and *P.aeruginosa* challenging medical practitioners and researchers to develop or discover new antimicrobial agents to combat these organism (Boucher 2010). Further, the pan-drug resistant in pathogenic bacteria has become more critical concern in public health gain urgent focus of research to find new and more effective antibiotic drugs (Walsh 2003, Talbot, Bradley et al. 2006). In spite of the urgent need the progress in development of new antibiotics is slow and the increase of new approved antibiotics has been steadily decreased in last few decades (Fischbach and Walsh 2009, Donadio, Maffioli et al. 2010). Therefore, to treat the infections, especially in hospitals where resistance immediately may become life threatening, the discovery of new antibacterial compounds is becoming more urgent (Shlaes, Projan et al. 2004). Bacilli are ubiquitous, endospore forming, rod shaped Gram positive bacteria that are of great importance due to their ability to produce broad spectrum of antibacterial peptides. These bacteria are extensively used in industrial processes for different enzyme production like proteases and also used as plant growth promoting bacteria (Posada, Alvarez et al. 2016). Their metabolites show antagonistic properties, so they are used against different pathogens (Al-Ajlani, Sheikh et al. 2007). *Bacillus subtilus* produce Biosurfectant iturin having significant antimicrobial activity (Kim, Ryu et al. 2010). Recently, biosurfectants are explored as alternatives to kill or inhibit growth of pathogenic bacteria (Dusane, Nancharaiah et al. 2010). Biosurfectant transform the exterior properties of bacteria and decrease adhesive potential (Walsh 2003). Morover, it has been discovered that biosurfectant produced by bacteria inhibit biofilm formation and interfere in bacterial cell to cell comunication (Irie, O’toole et al. 2005, Valle, Da Re et al. 2006).

Urinary tract infections (UTI) are the most prevalent infections occurs word wide especially in developing countries including Pakistan, where the chances of drug resistant is much higher due to injudicious use of antibiotics (Shah, Wasim et al. 2015). The majority of UTI is caused by *S.aureus, E.coli, K.neumomiae and P.aeruginosa* (Manikandan, Ganesapandian et al. 2011). Unfortunately these bacteria have been proved as MDR thus making its treatment ineffective (Shah, Wasim et al. 2015). These pathogenic bacteria also contribute to nosocomial infections and cause severe diseases in immunocompromized patients (Eguchi, Miyamoto et al. 2013). *A. beumannii* is gram negative bacterium ubiquitly found in environment and cause significant public health problems and is becoming increasingly important as a causative agent of nosocomial infection. The current study was therefore conducted to explore the indigenous soil bacterial flora for antibacterial potential that could kill or inhibit the causative agent of UTI and nosocomial infections. The indigenous soil bacteria *bacillus safensis* MK-12 or their antibacterial peptides might be employed to reduce or prevent the incidence of UTI and nosocomial infections.

## Materials and methods

### Soil sampling and bacterial isolates cultivation

Soil samples (9) were collected aseptically from different sites of district Kohat, Pakistan (33.5638° N, 71.4656° E) at the depth of 3-8 cm of the soil. The samples were serially diluted in sterile distilled water and 100 μl of each dilution was spread homogeneously with the help of sterilized swab or glass spreader on trypticase soya agar or/and nutrient agar (Oxide, UK) plates. The plates were incubated at 37 °C for 24-48 hours. A single bacterial colony was obtained by repeated sub culturing technique. Fresh pure culture was preserved in 20 % glycerol and stored at ×30 °C until required.

### Screening for antibacterial peptides production

All the 47 isolated bacterial isolates were first preliminarily screened for antibacterial activity by cross streaking method, agar overlaid method and/or spot inoculation method. In cross streaking method, individual soil isolate was streaked in the centre and test strains were streaked perpendicular of soil isolate (Peek, van Gelderen et al.). In agar overlay method, 20-24 hours old colony of soil isolate was inoculated on Muller hinton agar (MHA) plates and later on overlaid by 4-6 ml of semisolid medium with standardized 0.5 McFarland suspensions of test strains. In spot inoculation method the soil isolates were spot inoculated on MHA having pre uniform lawn of test bacteria. The clear zone of inhibition (ZOI) around the colonies was observed indicated antagonistic activity of the soil bacteria against test bacteria. Later on antibacterial activity of cell free supernatant (CFS) of soil isolates was determined against a set of test ATCC bacterial strain and clinical MDR bacterial strains (Table. 1). The clinical MDR strains were collected in another study from local tertiary care hospitals and resistant pattern was determined according to CLSI protocols. The secondary screening for antibacterial metabolites production was evaluated by agar well diffusion method or disk diffusion method on MHA plates.

**Table 1.** Optimized medium composition (component/ L)

| **Component** | **Quantity** |
| --- | --- |
| **Peptone** | 35 gm |
| **Sucrose** | 20 gm |
| **Yeast extract** | 8 gm |
| **Potassium di-hydrogen phosphate** | 2 gm |
| **Magnesium sulphate** | 0.045 gm |
| **Trace elements** | 9 mL |

**Table: 1.**
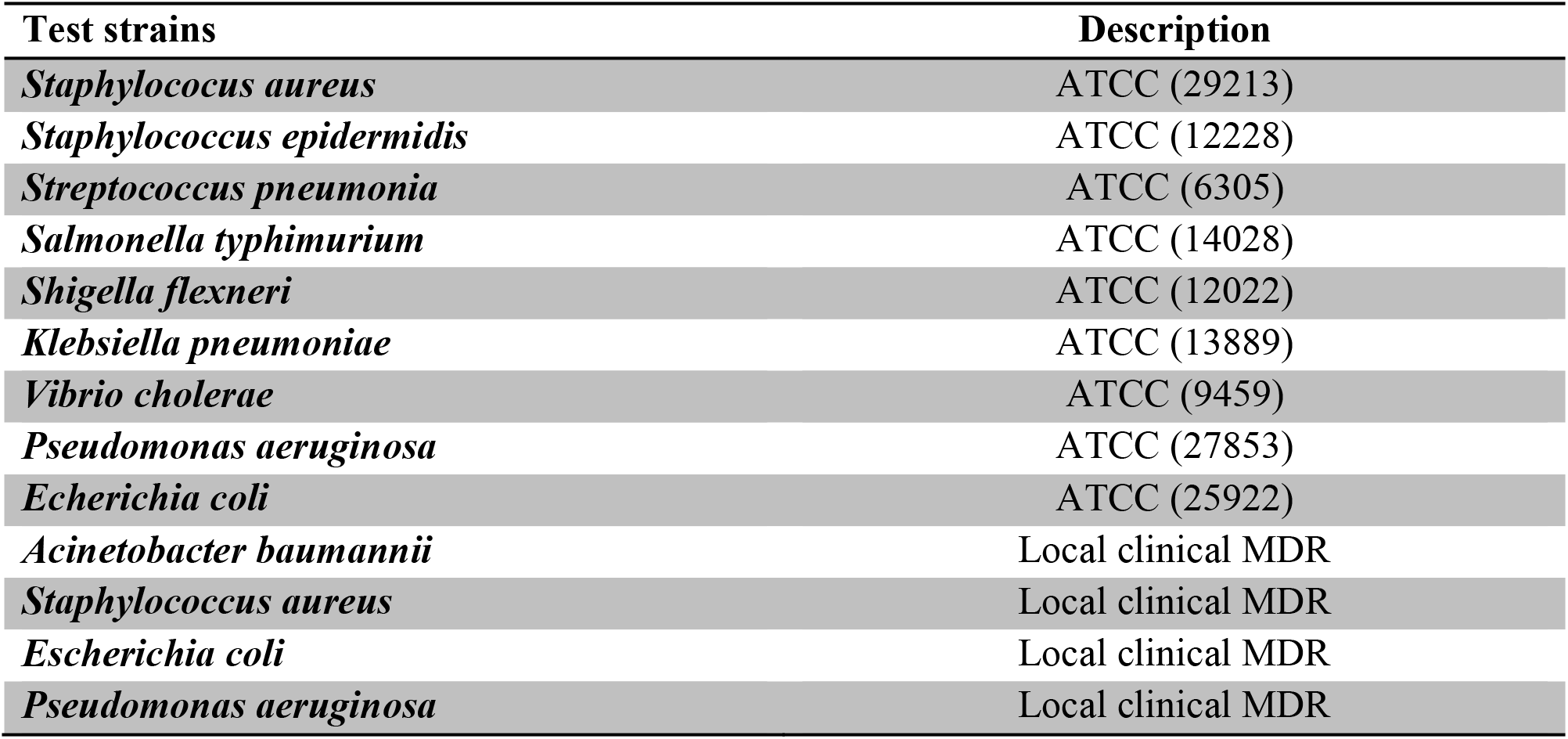

### Agar disk diffusion method

The antibacterial activities of CFS from soil bacterial isolates were checked by agar well diffusion method with a little modification as described by Kirby Bauer (Umer, Tekewe et al. 2013). The plates were kept at 4 °C for 1 hour to facilitate the diffusion of CFS in the agar medium and incubated at 37 °C for 24 hours. The selected standard procedure was applied in triplicate on the MHA for each soil bacterial crude extract.

### Agar well diffusion method

The wells (6 mm in diameter) were prepared by using sterile cork borer and standardize inoculum was uniformly streaked over MHA plate (Valgas, Souza et al. 2007). A volume of 70-80 μL of CFS obtained from different soil isolates was aseptically dispensed into each well. Sterilized water and vancomycine was used as negative control and positive control respectively.

After 24 hour of incubation, ZOI around the wells were measured and the experiment was performed in triplicate.

### Optimization of conditions for growth and antibacterial peptides production

Culture condition like temperature, medium, incubation time, pH and aeration were optimized for antibacterial peptides production as well as for growth. For temperature optimization, erlenmeyer flasks containing 250 ml TSB, 5 ml inoculums culture (equal to 1 McFarland) were inoculated and incubated at different temperatures such as 25, 30, 35 and 40 °C for 48 hours. The antibacterial activity was determined by agar well diffusion assay and bacterial biomass was estimated by spectrophotometer. For culturing medium optimization the producer isolate was cultured in different media i-e Luria-bertani broth, BHI broth, Nutrient broth, TSB and OPT medium with a little modification (Akpa, Jacques et al. 2001). All cultures were incubated at 37 °C for 48 hours and CFS was checked for antibacterial activity against test strain by agar well diffusion method. The effect of incubation time for optimum production of antibacterial metabolites was evaluated by inoculating 5 ml standardize inaculom (equal to 1 McFarland standard) of soil bacterial isolate in erlenmeyer flasks containing 250 ml medium incubated at 37 °C. Aliquots were collected at different time intervals (each 12 hrs for 96 hours) and antibacterial activity of CFS was determined against test strain by agar well diffusion assay. To evaluate the impact of pH on production of antibacterial metabolites, erlenmeyer flasks containing 50 ml TSB with 1 ml standardize inoclum of soil bacteria, were calibrated at different pH as 6, 7, 8 and 9. The flasks were maintained at 37 °C for 48 hours. After incubation antibacterial activity of CFS was determined against test strain by agar well diffusion assay. The effect of aeration on growth and antibacterial metabolites production was evaluated by culturing the producer isolate in shaker culture at various revolutions per minutes (rpm) and static culture for 48 hours. The growth of soil bacterial isolate was determined as O.D at 600 nm by using spectrophotometer at different pH, temperature, incubation time, medium and aeration.

### Antibacterial peptides production and partial purification

Antibacterial exhibiting soil bacteria was cultured up to late lag phase and then inoculated in erlenmeyer flasks containing 250 ml TSB, incubated at 37 °C for 48 hours in a shaking incubator. Subsequently, the culture was centrifuged at 4000 rpm, at 4 °C for 20 minutes and supernatant was collected. In order to remove all the cells and contaminants supernatant was filtered through 0.45 μm pore size filter paper. The pH of CFS was neutralized by adding 1M NaOH and 10 *%* HCl (Muriana and Klaenhammer 1991). To obtain optimum precipitation of antibacterial metabolites ammonium sulfate was added to CFS with continuous stirring at 4 °C till the level of saturation. The saturated CFS was incubated at 4 °C for 18 hours. Ammonium sulfate precipitates were separated by centrifugation at 4000 rpm, for 1 hour, at 4 °C. The supernatant and precipitates were obtained separately in sterilized test tubes. The pellet was dissolved in 50 ml of 25 mM sodium phosphate buffer (pH 7) and referred as crude antibacterial substances. Antibacterial activity was determined by agar well diffusion assay and ZOI was measured in mm.

### Determination of minimum inhibitory concentration (MIC)

Minimum inhibitory concentration of antibacterial peptides produced by *B.safensis* MK-12 was determined by broth 2 fold serial dilution technique. Trypticase soya broth (TSB) of 1 ml was poured in sterile screw caped test tubes from 1-10 and 2 ml TSB was poured into test tube no.11 (negative control). A volume of 1 ml partially purified antibacterial peptides was added into test tube 1 and 2 ml to test tube no. 12 (positive control). Initially 1 ml of homogenize solution was transferred from tube 1 to 2 and similarly all tubes were serially diluted except positive and negative control and incubated at 37 °C for 18-24 hours. The lowest concentration of antibacterial peptides that completely inhibits growth of tested strains was defined as MIC (Hoelzer, Cummings et al. 2011)

### Characterization of antibacterial peptides

To estimate molecular weight of antibacterial peptides the partially purified antibacterial peptides were diffused via dialysis membrane (pore size <20 kda moleculr weight). Partially purified antibacterial peptides were precipitated with 1 ml ammonium and were roofed in dialysis membrane in a beaker containing sterilized distilled water. All the setup was kept on agitator for 18 hours at 25 °C. The antibacterial activity of dialyzed antibacterial peptides was determined by agar well diffusion assay. To confirm the nature of antibacterial peptides, the partially purified antibacterial peptides were treated with proteinase K and subsequently antagonistic activity was evaluated by agar well diffusion method. Temperature stability of antibacterial peptides was evaluated by heating the CFS from 20 to 121 °C for 20 minutes and antibacterial activity was determined by agar well diffusion method.

### Haemolytic activity

Haemolytic activity of antibacterial peptides was carried out by blood agar plate technique. The blood agar plates were prepared by adding 5% of human blood to agar base. Sterile gel borer was used and wells were punched on blood agar plate. The ethyl acetate extract of 1000 μg/mL was prepared and 70 μl was aseptically dispensed in the well. The blood agar plates were incubated at 37 °C for 24 hours.

### Antibacterial activity of partially purified antibacterial peptides

Antibacterial activity of ammonium sulfate precipitates of antibacterial peptides was assayed against 4 local MDR clinical bacterial strains including *P.aeruginosa, A.beumanii E. coli and S.aureus*. The test strains standard inoculom (equal to 0.5 McFarland) was uniformly streaked on the surface of solidified MHA plate. Wells were made and 70 μl of partially purified antibacterial peptides were dispensed and incubated at 4 °C for 2-3 hours to facilitate the diffusion of bioactive peptides into the agar and later on, plates were incubated at 37 °C and ZOI was measured after 24 hours. Each experiment was performed in triplicate.

### Identification of soil isolate *Bacilus safensis* MK-12

Preliminarily the soil isolate MK-12 was identified by Colony morphology, Gram reaction and biochemical profiling according to bergeys manual of bacteriology. Subsequently the soil isolate was identified by 16S rRNA gene sequencing. Genomic DNA was obtained by phenole/chloroform extraction method with a little modification (He 2011) and 16S rRNA was amplified by using oligonucleotide primers 9F (5-GAGTTTGATCCTGGCTCAG-3) and 1510R (5-GGCTACCTTGTTACGA-3) as forward and reverse primers in PCR. In PCR initial denaturation was performed at 94 °C for 2 minutes, annealing temperature was maintain at 52 °C for 90 seconds and extension time 2 minutes at 72 °C, and then the reaction was repeat for 35 cycles, follwed by final extention for 10 minutes at 72 °C. The amplified PCR product was separted on 1 % agarose gel and recovered by PCR purification kit (Roche, Germany). The purified product was sequenced by di-deoxy chain termination method (Macrogen, Korea) and the result was BLASTn in NCBI (https://www.ncbi.nlm.nich.-gov/BLAST/) and Eztaxon (https://www.ezbiocloud.net/identify). The 16S rRNA gene sequences of the related strains were retrieved from Ezbioloud to infer the evolutionary relationship of *Bacilus safensis* MK-4. The 16S rRNA sequence alignment and treming were performed by using Clustal omega (https://www.ebi.ac.uk/Tools/msa/clustalo/) and BioEdit respectively. Phylogenetic tree was constructed based on neighbor joining method by using a computational tool MEGA-7 (Kumar, Stecher et al. 2016). The reliability of the phylogenetic trees was determined by using boot strap technique on 1000 replicates.

### Statistical analysis

All the data obtain from secondary screening was analyzed by one way ANOVA. The level of significance was determined using SPSS version 15 and the results having P value <0.05 were defined to be significant.

## RESULTS

### Soil sampling and plate count

A total of 9 soil samples were collected and 47 bacterial isolates were purified. The sampling sites including hilly area, lawn, garden, industrial area, open cultivated field, river bank, stream bank, rhizosphere, road side, pounds surrounding area. Plate count on TSA ranged from 4 ×10^5^ to 7 × 10^5^ CFU/g of soil samples. Considerably lower CFU count (4 ×10^5^/g) was calculated for the soil sample collected from mill effluent soil sample Kohat (Table 2).

### Antibacterial activity

Among the total of 47 soil isolates 12 isolates (25.53 %) exhibited antagonistic activities against Gram positive ATCC bacterial strains or/and Gram negative bacterial strains. Among these 12 AEB, three isolates (25%) which were designated as MK-9, MK-10 and MK-32, showed antibacterial activity only against Gram positive bacterial strains while nine isolates (75 %), were active against both Gram positive and Gram negative bacterial strains (data not shown here). One isolate *B.safensis* MK-12, exhibited strong antibacterial activity against Gram positive, Gram negative ATCC as well as against MDR clinical strains.

**Fig: 1.**
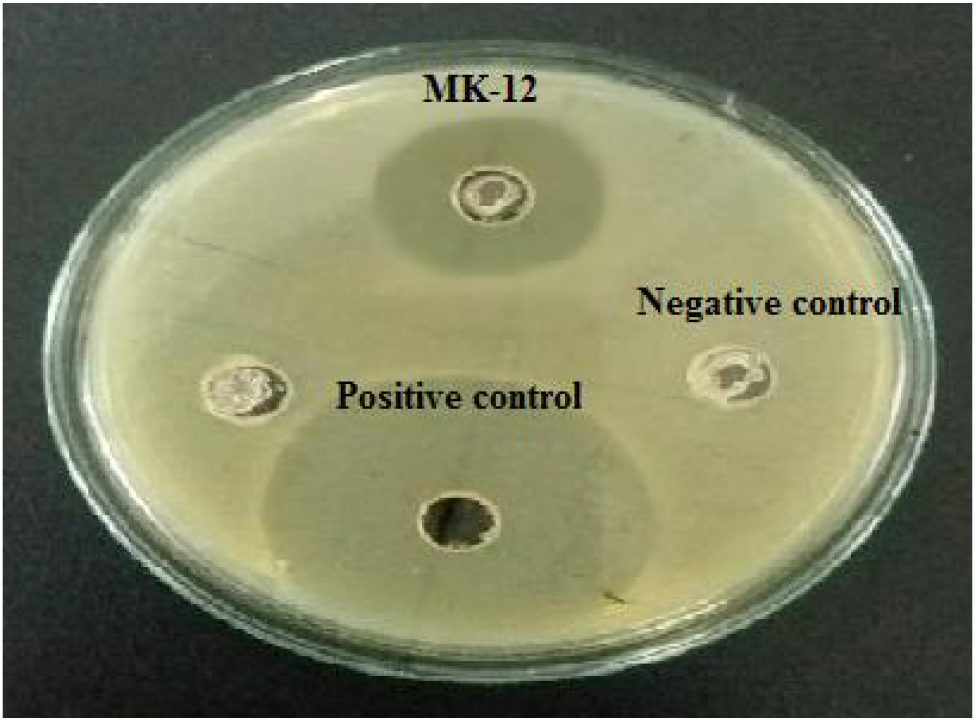
Showing antibacterial activity of partially purified peptides produced by *B.safensis* MK-12 against *S.aureus*.

There was noteworthy difference among antibacterial activity of soil isolates against tested bacterial strains. The antibacterial peptides (CFS) produced by soils isolate *B.safensis* MK-12 exhibited highest antibacterial activity (18+0.5 mm) against *S. typhimurium* (14028) as compared to other tested strains, while the tested strain *K. pnemoniae* (13889) showed higher resistant towards the bioactive peptides produced by *B. safensis* MK-12 and 15 mm ZOI was observed (Fig:2).

**Fig: 2.**
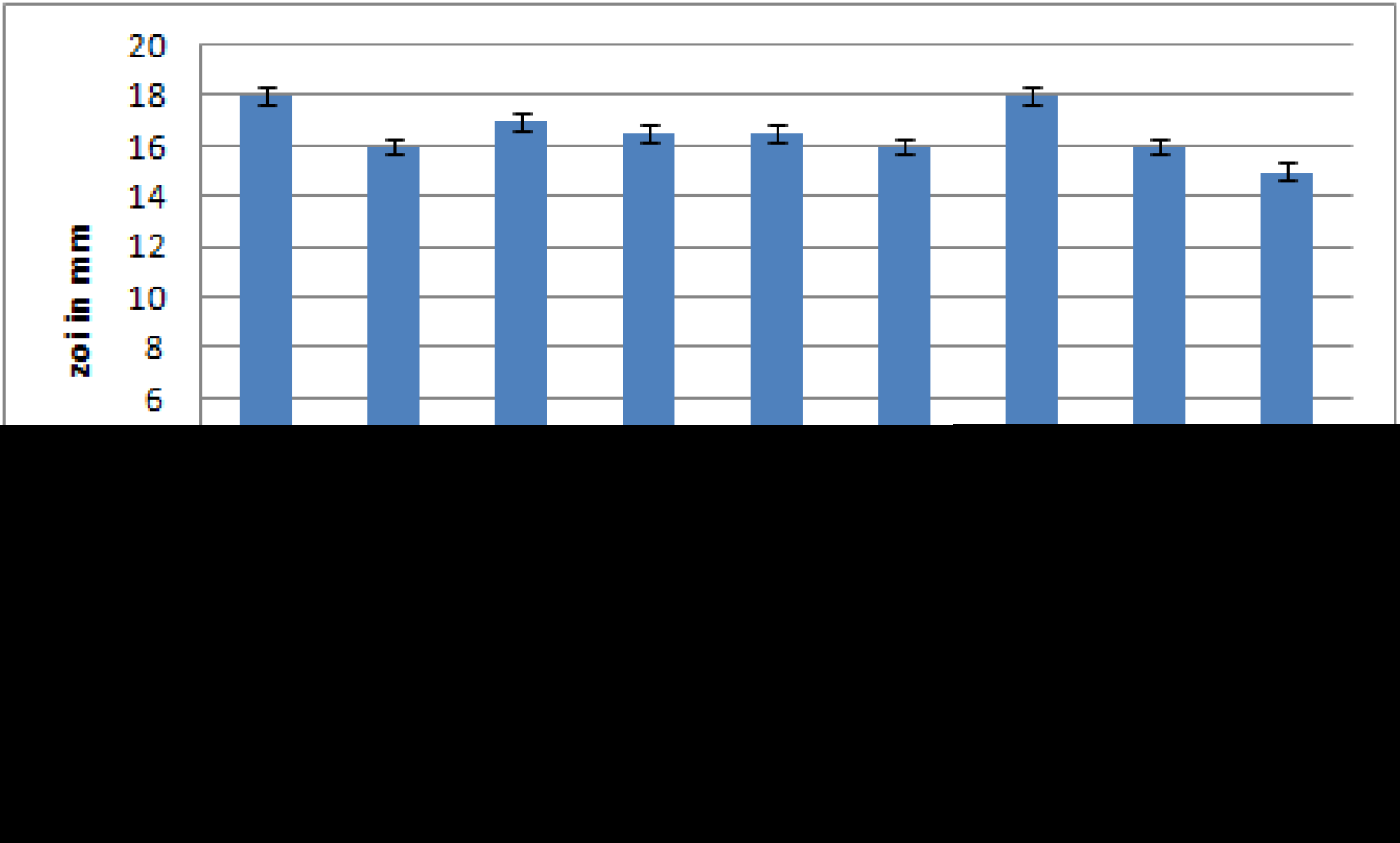
The antibacterial activity of *B.safensis* MK-12 against test ATCC bacterial strains

### Optimization of growth and antibacterial metabolites

Culturing medium, incubation time, pH and temperature were optimized for maximum growth and antibacterial metabolites production. Maximum antibacterial activity of CFS was achieved at 30 °C with 18 mm ZOI followed by 35 °C with inhibitory zone of 17 mm. The minimum antibacterial activity was exhibited at 45 °C, indicating that the soil isolate *B.safensis* MK-12 produce highest amount of antibacterial metabolites at 30 °C (Fig. 3). These results are in line with the studies of Kim *et al.*, 2006 in which *Enterococcus faecium* RZS C5 produced high amount of antibacterial metabolites between 25 to 35 °C than at 20 °C. Temperature has a profound effect on antibacterial metabolites production and denatures it to some extent. The current study result showed that nitrogen rich media i-e BHI and TSB enhanced the growth and antibacterial metabolites production as compared to simple nutrient media. The CFS obtained from optimized medium showed 19 mm ZOI, while CFS obtained from BHI and TSB medium exhibited 18 mm ZOI than the nutrient broth (14 mm), High amount of biomass was obtained by cultivating the producing isolate in optimized (opt) medium, TSB and BHI medium respectively, indicating that nutrients have considerable effect on antibacterial metabolites production. The BHI and TSB culture media is rich in carbon and nitrogen source as compare to nutrient broth as previously demonstrated by (Morais and Surárez 2016).

**Fig: 3.**
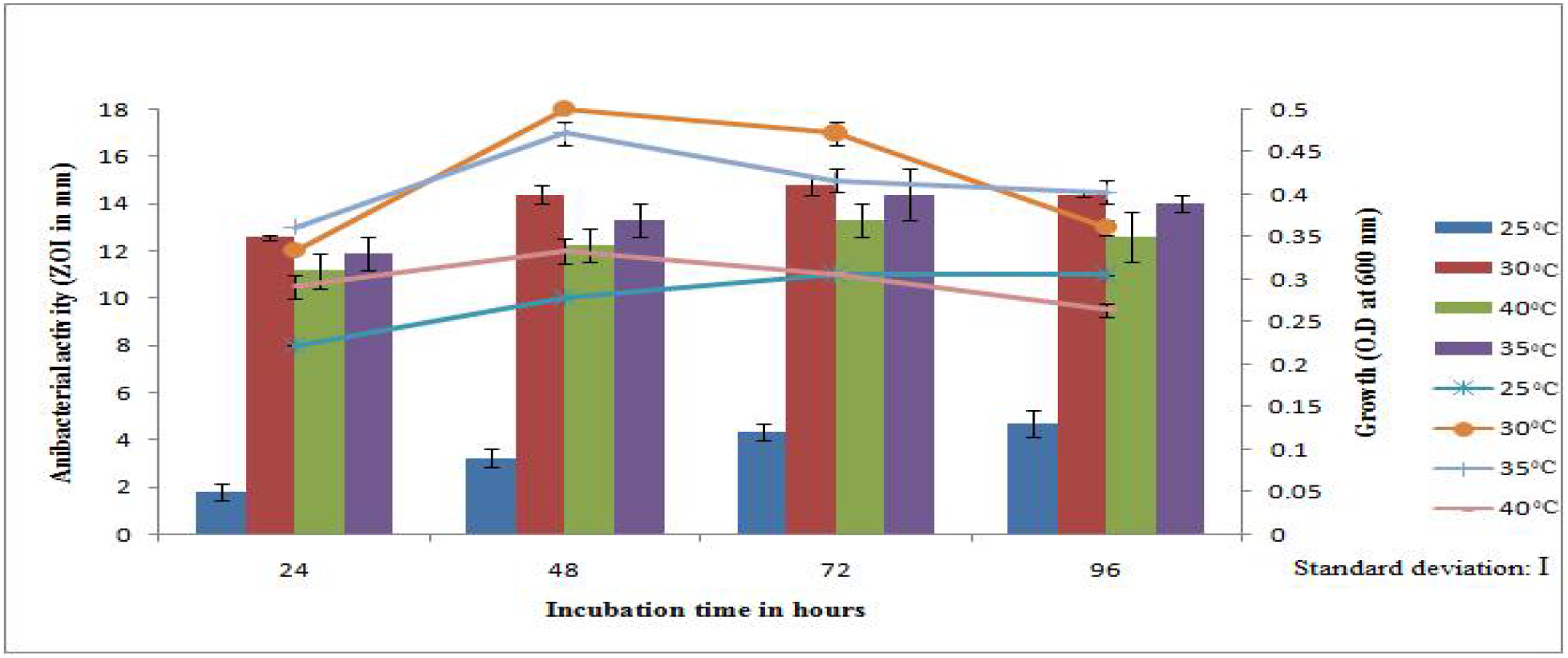
Effect of temperature and incubation time on growth and antibacterial activity of *B. safensis* MK-12 against *S.aureus*

The effect of pH on antibacterial peptides production was evaluated and exhibited maximum ZOI against test strains at pH 8, while comparatively low ZOI was recorded at 6, 7 and 9 pH indicating that pH 8 is optimum for antibacterial peptides production. Bacteria produce maximum antibacterial peptides at their physiological pH as declared by Parente and Ricciardi 1994 during enterocin from enteric bacteria and pediocin production. The possible reason of this phenomenon is the favorable pH for the growth of bacteria that facilitate the production of antibacterial peptides. In the current study maximum production time was 48 hours after that a decline in the ZOI was observed (Fig. 4).

**Fig: 4.**
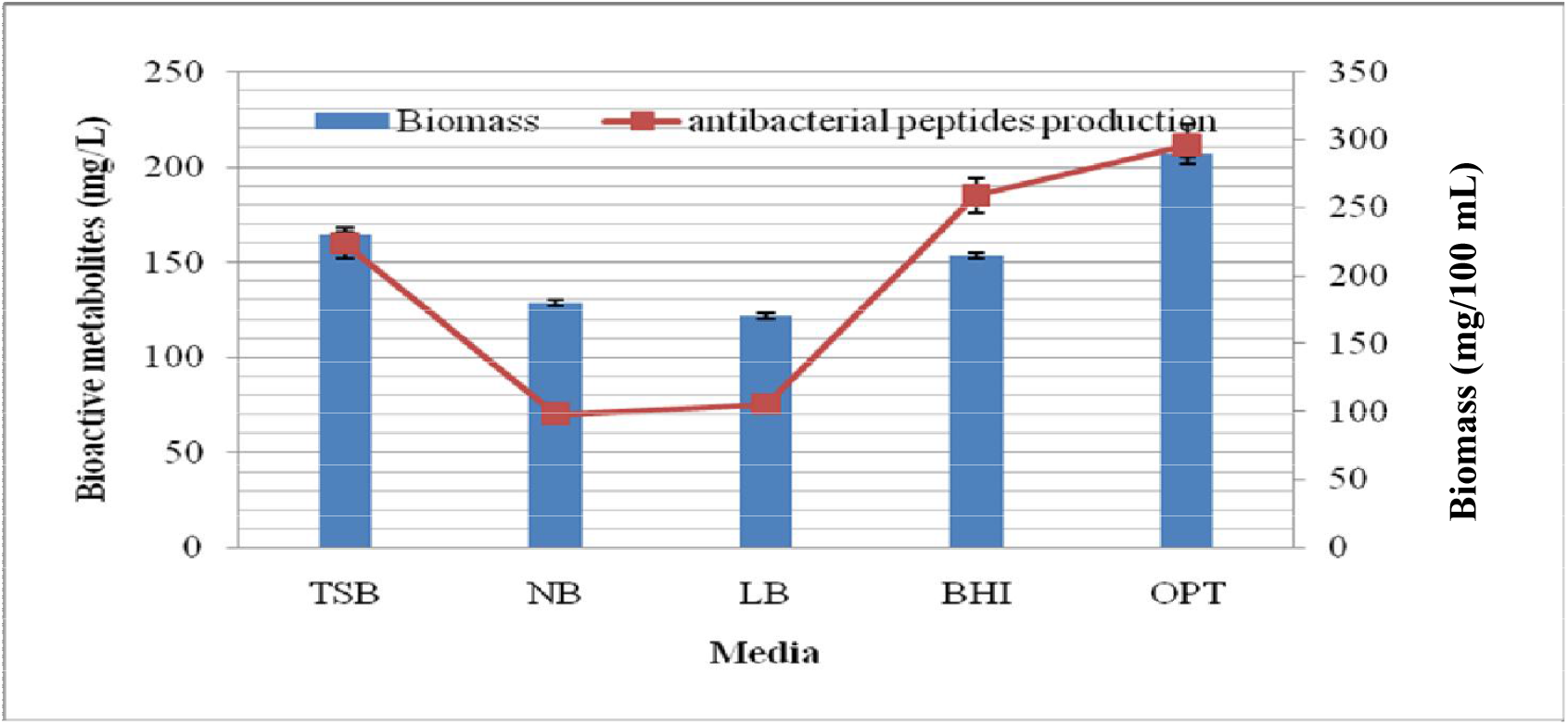
Antibacterial peptides production by *B.safensis* MK-12 grown on various culture media. Trypticase soya broth (TSB), Nutrients broth (NB), Luria bretani (LB), Brain heart infusion (BHI) and Optimized (Optm) broth medium.

**Fig: 2.**
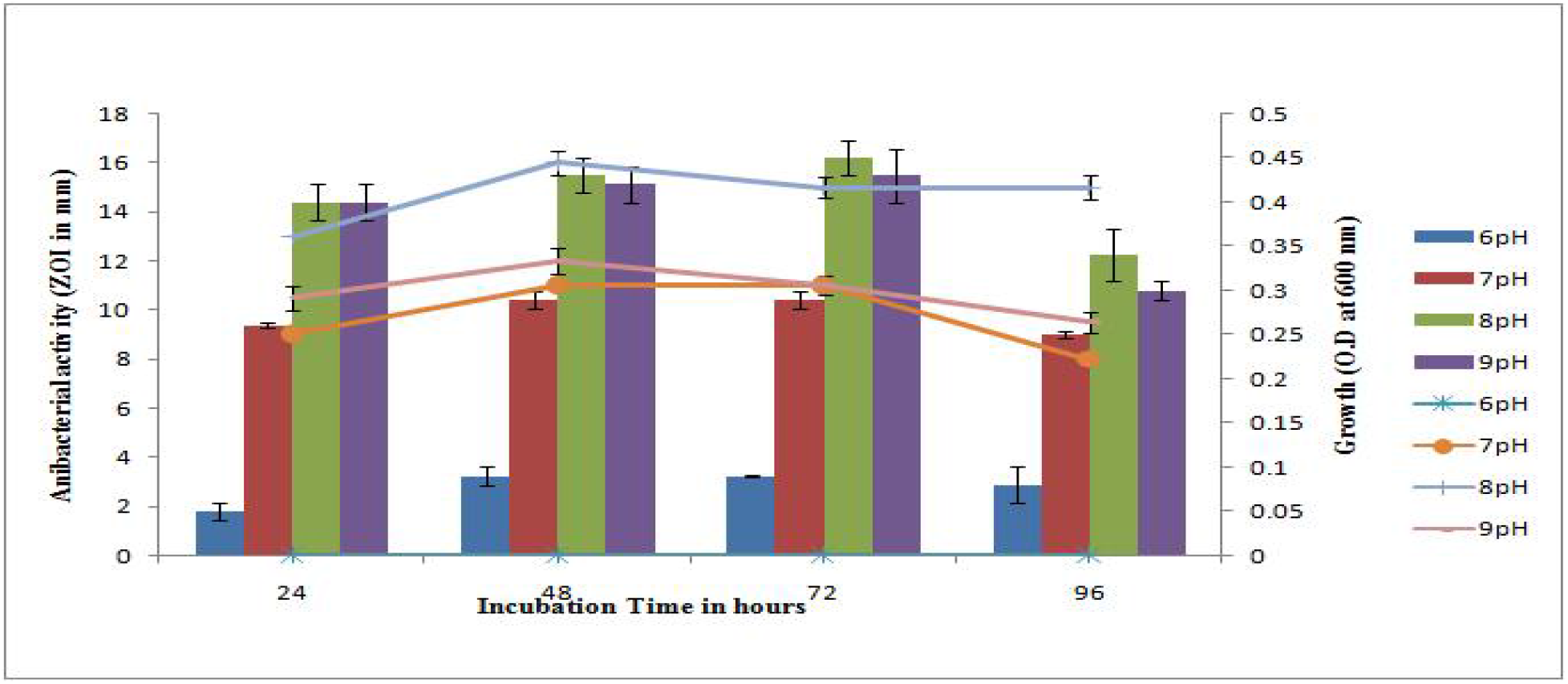
Effect of pH and Incubation time on growth and antibacterial activity of *B.safensis* MK-12 against *S.aureus*

### Characteristic and antibacterial activity of partially purified antibacterial peptides

The partially purified antibacterial peptides exhibited strong antibacterial activity as compared to CFS. The molecular weight of the antibacterial peptid was estimated less than 20 kDa. The ammonium sulfate precipitate treated with proteinase K lost their antibacterial activity validated that the antibacterial metabolites are protein in nature. When the CFS was heated from 20 to 121 °C for 20 minutes the antibacterial activity against test strains decreased at 45 to 100 °C and abolished when heated up to 121 °C. The sensitivity of antibacterial metabolites to proteinase K and high temperature confirm that it is protein in nature. Haemolytic activity of partially purified antibacterial peptides was evaluated against human blood and no activity was observed against human erythrocytes.

### Determination of MIC

MIC of partially purified antibacterial peptides ranged from 2.64 mg/mL to to 3.45 mg/mL against Gram positive ATCC bacterial strians. While against Gram negative ATCC bacterial strains the MIC ranged from 2.75 mg/mL to 3.85 mg/mL (Table 3).

**Table: 3.**
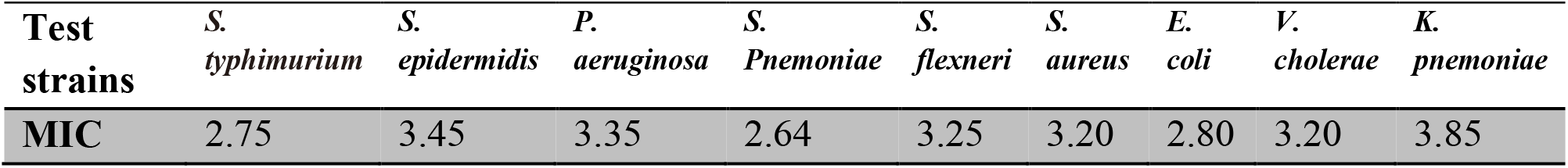
MIC of antibacterial peptides produced by *B.safensis* MK-12 (mg/mL).

| Test strains | *S. typhimurium* | *S. epidermidis* | *P. aeruginosa* | *S. Pnemoniae* | *S. flexneri* | *S. aureus* | *E. coli* | *V. cholerae* | *K. pnemoniae* |
| --- | --- | --- | --- | --- | --- | --- | --- | --- | --- |
| MIC | 2.75 | 3.45 | 3.35 | 2.64 | 3.25 | 3.20 | 2.80 | 3.20 | 3.85 |

### Morphological, biochemical and molecular identification of AEB

The soil isolate *B.safensis* MK-12 was characterized morphologically, biochemically and molecularly by 16Sr RNA gene sequencing. Result of the morphological characteristics of the isolate MK-12 revealed that the growth was excellent on TSA. Colony morphology of *B.safensis* MK-12 was irregular in shape, with undulant margin, raised elevation, light yellow in color and opaque opacity. Cell morphology showed that it is Gram positive, rod shaped, motile and spore forming bacterium. Biochemical results indicate that *B.safensis* MK-12 is positive for ureas, catalase, oxidase production and MR test while negative for indole, citrate and VP test. The 16S rRNA gene sequence analysis of MK-12 revealed a high level of similarity (98%) to the sequence of *B. safensis* and *B. pumilus*.

**Fig:**
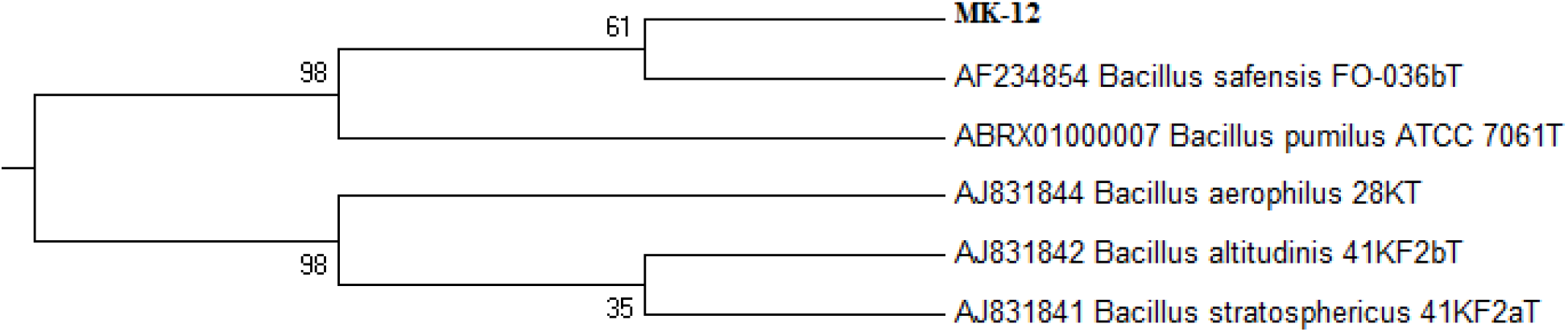
Phylogenetic tree constructed showing evolutionary distance based on 16S rRNA gene sequence by using neighbor joining method, showing the phylogenetic residency of isolate *B. safensis* MK-12 and other related members of the genus *Bacillus*. Bootstrap percentages (based on 1000 replicate) are indicated at each branch nodes.

## Discussion

Currently, the prevalence of multi-drug resistant bacteria is increasing and conciliation the treatment of an increasing number of infectious diseases. Thus, due to the burden for high frequency of MDR pathogenic bacteria in the world there is an urgent need for searching new drugs which are effective against current antibiotic resistant pathogens. In the current study, the soil samples were collected from diverse localities for antibacterial metabolites producing bacteria. A total of twelve (25.53%) out of 47 bacterial isolates were exhibited antibacterial activity in primary screening. This result indicate high no. of AEB than 21.88%, less than 42% and approximately equal percentage 26.7% from previous reports (Abo-Shadi, Sidkey et al. 2010, Al-Ajlani and Hasnain 2010, Bizuye, Moges et al. 2013).

In the current study, the ranges of ZOI of CFS against test bacteria were 15-18 mm. These results are in line with the study (Gurung, Sherpa et al. 2009) where 0-18 mm ZOI was recorded for partially purified metabolites from soil bacteria against selected test organisms. The CFS was further partially purified and antibacterial activity was evaluated against MDR *S.aureus*, *E.coli, A.beumanii* and *P.aeruginosa* as well as against ATCC strains. The ZOI against MDR was ranged from 14-18 mm which was found to be higher when compared to Yucel and Yemac’s study results (Yücel and Yamac 2010). The results of the present study are fascinating and encouraging because the partially purified metabolites from the soil isolates *B.safensis* MK-12 exhibited promising antibacterial activity against both ATCC and MDR clinical pathogens. According to the current study result, vancomycin showed 24 mm ZOI against MDR local clinical strains which is higher as to partially purified metabolites from the soil isolates. But it is expected that further complete purification of the antibacterial peptides will exhibits good ZOI as showed by vancomycin after complete purification. Therefore, further purification process is important to get pure antibacterial metabolites for the application of treatment of various pathogenic bacteria.

Maximum antibacterial activity CFS from soil isolate *B.safensis* MK-12 against test strains was obtained at 30 °C with 18.6 mm ZOI followed by 37 °C with inhibitory zone of 18 mm. The minimum antibacterial activity of CFS was recorded at 45 °C, indicating that the producer isolate can produce highest amount of antibacterial metabolites at 30 °C. The lesser inhibitory zone was noted at lower and higher temperature than 30 °C (Fig. 3). These results are in line with the other study in which *Lactobacillus spp*. yield highest amount of bacitracine at 25 and 30 °C (Malheiros, Sant’Anna et al. 2015). Temperature significantly affects the antibacterial metabolites production and may inactivate it to some extent. In previous studies, *Enterococcus faecium* RZSC-5 produced higher antibacterial peptide between 25-35 °C than at 20 °C (Leroy and De Vuyst 2002). The nitrogen rich medium (BHI) facilitate the growth and antibacterial activity of producer bacteria compared to simple nutrient medium. The ZOI of CFS taken from BHI was higher (25 mm) than the nutrient broth (15 mm), indicating the nutrients have significant effect on production of antibacterial metabolites production. The BHI and TSA is a source rich in nitrogen and carbon as against nutrient broth as discussed by (Leroy and De Vuyst 2002). Multi drug resistant Gram positive strains such as *S. aureus* are often causing UTI and nosocmial infections become resistant to antibiotics by modifying its teichoic acid in cell wall with D-alanine at the result increased MIC. The result of current study is worth observing that the MIC against MDR strains not increased and were comparable to other antibacterial metabolites produced by *bacillus spp*. The MIC results varied among the tested strains against Gram positive and Gram negative bacteria. These results were in agreement with the study of Sibanda et al (Sibanda, Mabinya et al. 2010). The antibacterial ethyl acetate extract of *B.safensis* MK-12 is resistant to temperature and did not exhibit hemolytic activity when tested against human blood cells which could be considered as safe for human use.

In conclusion the results of current study demonstrate that northern region soil of Pakistan is rich with respect to antibacterial compounds producing bacteria from which novel bioactive compounds can be elicited. The AEB or their bioactive metabolites could be used as a biocontrol to reduce the pathogenic bacteria in environment. Moreover these bioactive compounds may be good candidate for the development of new therapeutic agents.

